# Rapid Nanopore cloud-based monitoring and analysis of commercial aquaculture microbiomes

**DOI:** 10.1101/2024.08.07.607111

**Authors:** James P. B. Strutt, Irina Bessarab, Jamine Goh Ying Min, Hana Marican, Stephen Summers

## Abstract

Aquaculture stands as a crucial component of global food security and sustainable development, yet it faces challenges in disease management and ecological balance. Here, we present a novel approach using rapid nanopore sequencing, cloud-based monitoring and analysis for aquaculture microbiomes. Our study aimed to perform untargeted, agnostic biological monitoring of a commercial aquaculture facility, emphasizing rapidity, specificity, and sensitivity. We employed Oxford Nanopore Technologies’ MinION sequencer with an optimised rapid sequencing protocol, enabling on-site operation by facility staff. Three separate sampling efforts resulting in thirteen sequencing runs were conducted, revealing a representative microbiome baseline across aquaculture system components within a 24-hour timeframe. Our results demonstrated the feasibility of rapid monitoring and analysis of nitrogen-associated organisms, essential for same-day water quality management and infection event detection. Notably, Moving Bed Biofilm Reactor (MBBR media) or BioDiscs exhibited the highest diversity and abundance of nitrogen-associated organisms, confirming a pivotal role in nitrification processes. Critically, our approach addressed challenges in metagenomic sample purity and false positives, offering insights for future refinement and application. Our findings underscore the potential of rapid sequencing technologies in enhancing aquaculture management and sustainability. This approach holds promise for mitigating disease outbreaks, optimizing productivity, and advancing ecological balance in aquaculture systems.

## Introduction

Aquaculture is vital globally as a sustainable solution to meet escalating food demands. With wild fish stocks declining, aquaculture ensures a consistent protein source while conserving ocean ecosystems.^1^ It contributes to food security, reduces pressure on overfished species, and supports regional economic growth, making it indispensable for a nourished and balanced future. Aquaculture plays a pivotal role in achieving several United Nations Sustainable Development Goals (SDGs) such as SDGs 2 (Zero Hunger), 8 (Decent Work and Economic Growth), and 14 (Life Below Water). It aligns with SDG 2 by bolstering food production and enhancing nutritional accessibility. By reducing pressure on overexploited marine resources, it supports SDG 14 and helps restore aquatic ecosystems. Furthermore, aquaculture’s economic contribution addresses SDG 8 and promotes resilient livelihoods, especially in developing nations. Embedded within these sustainable goals is the recognition of the critical link between aquatic ecosystem health and human well-being. The assessment of the aquaculture microbiome emerges as a catalyst in achieving these objectives. Understanding the intricate relationships between aquatic microbiomes, fish health, and environmental stability is essential for progressing toward these global targets. By harnessing the power of microbiome research, we can unlock innovative solutions that propel us closer to a more resilient, nourished, and sustainable future.

Understanding the microbial world in aquaculture is crucial to prevent disease outbreaks, ensuring healthy aquatic species, optimizing productivity, and maintaining ecological balance, thereby safeguarding the industry’s economic and ecological viability. The importance for timely detection of a disease outbreak within an aquaculture setting is important on multiple levels. In aquaculture, marine diseases can spread rapidly and be highly problematic, due to crowded conditions of the stock organisms.^2^ This can result in the loss of the entire harvest. This includes cases of the cultured seafood having an unpleasant taste as a result of poor health or appearing unappetizing, as with cases of bitter crab disease, it can lose all commercial value. We see a similar impact with infected fish muscle tissue, resulting in higher processing costs and therefore lower revenue (ref?). Lafferty, *et al*^3^ gives examples of economically consequential marine diseases. 67 cases were reviewed with a breakdown ofd infections caused 18% by metazoans, 19% by protists, 34% by bacteria, and 25% by viruses. With regards to the hosts that were influenced by these economically consequential diseases, the most impacted were fish species with 49% of commercial species being impacted by disease, then molluscs at 28%, followed by crustaceans at 21%. Also noteworthy is that most of the studies were conducted in temperate and not tropical waters, suggesting that there is a paucity of understanding of these economically important diseases in the tropical environments.^3^

While culture-based methods play a major role in the identification of some pathogens, molecular methods, such as polymerase chain reaction (PCR) have increasingly been used for the molecular detection of animal diseases^4^. Prior work also demonstrated the use of whole genome sequencing approaches using Illumina MiSeq platform for detection of antibiotic resistance in pathogenic bacteria in hospitals within a day^5^. Moreover, whole genome sequencing on the Illumina MiSeq platform was used for monitoring non-typhoidal Salmonella as a foodborne pathogen in surface waters from *Salmonella enterica*^6^. One of the key impediments to the rollout of sequencing based diagnostic tests for genomic surveillance are the major costs required and the skill sets involved in analysis. As such, any solution to this problem needs to be affordable enough for the operator to consider using it as part of their daily operations. Simultaneously, a key requirement is that any downstream analyses of the synthesised data be easily interpretable by a non-specialist^7^.

### Relevance to Aquaculture Facility Management

As a means of systems management, DNA sequencing has been used for nitrogen cycle surveillance by examining the microbial community.^8^ This includes surveying anammox bacteria, ammonia oxidizing bacteria and nitrogen oxidizing bacteria. Typically, samples can be taken 3-5 times a week and processed once per week per the protocol of Andersen *et al.,*^9^. Their work suggests that DNA surveillance was at least an important supplement and, in many cases, outperformed sensor data for managing aquaculture systems. While the work undertaken by Andersen *et al.,*^9^ focused on examining the nitrogen cycle, a similar workflow could be applied to disease monitoring via the microbiome^9^ for various regions of any system, such as the host animal, biofiltration systems, or the water column. Using 16S rRNA gene amplicon sequencing, fish gut microbiomes were examined by Karlsen *et al.,*^10^. They observed that environmental DNA (eDNA) within animal feed acted as a confounding factor, which led to the enrichment of a few taxa, including *Mycoplasma* and the *Ruminococcaeae*. This feed signal is detected within the host microbiome, and therefore can result in inaccurate assessment due to the presence of this exogeneous source of DNA. This remains a consideration when assessing fish-related samples, especially if the fish are being sampled as part of the genomic surveillance approach^10^. In medical settings, similar metagenome approaches have been used to try and understand sample sterility of cell cultures in cell therapy manufacturing.^11^ However, this presents a relatively different problem than the current one as the starting sample should be comprised only of human sequences and will only present as positive if there is a contamination event. Nonetheless it is important to be able to detect these instances of contamination to a high level of sensitivity and similar techniques and levels of accuracy can be used to understand these aquaculture metagenomes^11^.

### Objective of the Project

The aims of this project were to develop a system for rapid biological monitoring of an aquaculture facility from an untargeted, unbiased perspective. Ideally, the solution to this would be rapid, with the entire process taking less than 24 hours to run. The tool should be specific and identify the contaminant species that might be causing deleterious effects on the fish or aquaculture stocks, as well as to be sensitive and detect low abundant organisms in addition to those that are present in high abundance. The analysis requires a one-day process as contaminant infectious agents can spread to the entirety of the system within a relatively short period of time. Therefore, catching these contamination events early allows for rapid and early husbandry and veterinary intervention that will provide healthier fish and minimize the requirement for lengthier veterinary treatment protocols.

The final component required by the technology is that it should be conductible without the requirement for a specialised microbiological or molecular biology laboratory. The solution should be feasible at the aquaculture facility itself. Staff can be trained to run these analyses in real time, as frequently as required, and as financial considerations permit. This is important because some facilities may be remote and lack sufficient infrastructure to rapidly sample and quarantine infected stock samples. Therefore, sampling and analysing on-site can mitigate this deficiency in infrastructure, as well as enabling rapid veterinary intervention. The tool that allows us to adequately address these requirements is the Oxford Nanopore Technologies MinION, a third-generation portable sequencer. In conjunction with the rapid sequencing kit, we can generate long DNA reads that allow for high quality identification of these aquatic metagenomes, such as has field-based monitoring of biothreats using simulated metagenomes.^12^

The scope of the work was to identify a microbiome baseline present within the aquaculture system waste (sump, drum tank and BioDiscs) and develop an approach to track the metagenomes of tropical fish farms in near real-time and develop mitigation strategies against repeated disease outbreak events. We partnered with Singapore Aquaculture Technologies (SAT) Pte Ltd, a Singaporean based aquaculture facility that specialises in the growth and monitoring of Asian Sea Bass to address this problem. Our project aimed to optimise for long read sequencing in the aquaculture settings. We performed 3 separate sampling efforts with 13 sequencing runs over a 6-month period for methods development occurring in parallel, demonstrating we can detect a representative baseline of the microorganisms thought to be prevalent in the aquaculture facilities, in a 24-hour time frame, which allows for rapid response.

## Materials and Methods

### 2.1. Sample Collection and Storage

All samples were collected from a free-floating recirculating aquaculture farm, positioned in the Serangoon Harbour, Singapore (1.397N, 103.935E) and owned and operated by Singapore Aquaculture Technologies (SAT) Pte Ltd. The farm is comprised of ten open air barramundi holding tanks fed from a drum filter and sump tank system, house under a roof powered by solar panels. Samples for DNA extraction were sourced primarily from the filtration and waste systems of the aquaculture facility, which are the drum tanks and the sump. The facility rears Barramundi *spp*., alongside Red Snapper and Grouper. MBBR media (BioDiscs), used for water filtration by providing nitrogen processing biofilm friendly growth conditions, were collected for assessment from the filtration systems. After collection, samples were immediately stored in falcon tubes and processed as described below.

### 2.2. DNA Extraction

Samples from the sump and drum tank were prepared using a weight range of 100-150mg of particulate flocculation per extraction. For the samples from MBBR, a quarter of a BioDisc chip was used for each individual extraction. The DNeasy PowerSoil Pro Kit (Cat#47014, QIAGEN) was utilized for the DNA extractions according to the manufacturer protocol except that the fraction of the BioDisk was combined with the Powerbead Pro beads in 15 mL centrifuge tube for the bead beating step. Samples were vortexed in Vortex-Genie 2.0 mixer at 2/3 speed for 5 min. Following extraction, the DNA concentration was quantified with a Qubit dsDNA Broad Range Assay Kit (Cat#Q32850, ThermoFisher). Where required, additional purification step was performed using the the Zymo Genomic DNA Clean and Concentrator Kit (Cat#D4064).

### 2.3. DNA Sequencing

DNA libraries were prepared using the Rapid Sequencing Kit (SQK-RAD004, Oxford Nanopore Technologies (ONT)) or the Rapid Barcoding Kit (SQK-RBK004, ONT). The MinION flow cell version 9.4.1 was employed for sequencing. Samples were sequenced on Min MK1C sequencer (ONT) for 3 hours to access a rapid processing timeline. Flow cells were re-used after removing sequencing libraries with flowcell wash kit (WSH002, ONT). Software versions were as follows: *MinKNOW* v21.11.7, *MinKNOW* Core v4.5.4, *Bream* v6.3.5, *Guppy* v5.1.13.

### 2.4. Bioinformatics Pipeline, and Data Analysis

A custom pipeline was employed that accepted the base called FASTQ files as inputs. Adapter sequences were identified and trimmed from the individual reads using *Porechop* 0.2.4 (https://github.com/rrwick/Porechop). The metagenomic classifier, *Centrifuge*^13^, was used to classify reads, allowing identification of the metagenome from the samples. A comprehensive database amalgamating fungal, viral, and bacterial RefSeq genomes sourced from NCBI served for species-level classification, the bacterial component of the database was reduced to control for entries with high numbers of genomes, reducing the database size to 1/6 prior to addition of viral genomes and fungal genomes (including incomplete genome entries). Python was harnessed to wrap the aforementioned tools and databases into an integrated pipeline. Coverage estimates were conducted employing *<u>Minimap2</u>*^14,15^. *Minimap2* was used to compare sequences against the SILVA NR99^16^, both large and small subunits, to evaluate species alignment. In Figure 3, depicted are the relative abundance of aligned bases as calculated by taking hitLength from centrifuge for each taxon and dividing by hitLength sum to get aligned bases for a given sample for nitrogen associated organisms from these samples.

## Results

The developed workflow described in Figure 1A shows the three sampling locations from the aquaculture fish barge. In Figure 1B, the sample preparation, library preparation, cloud-based data from the barge and subsequent analysis are depicted. This workflow demonstrated that the microbiome of the aquaculture could be monitored in real-time using long-read sequencing aboard the barge, demonstrating at a basic level that this protocol can be completed without having to transfer DNA samples to a land-side laboratory.

**Figure 1.**
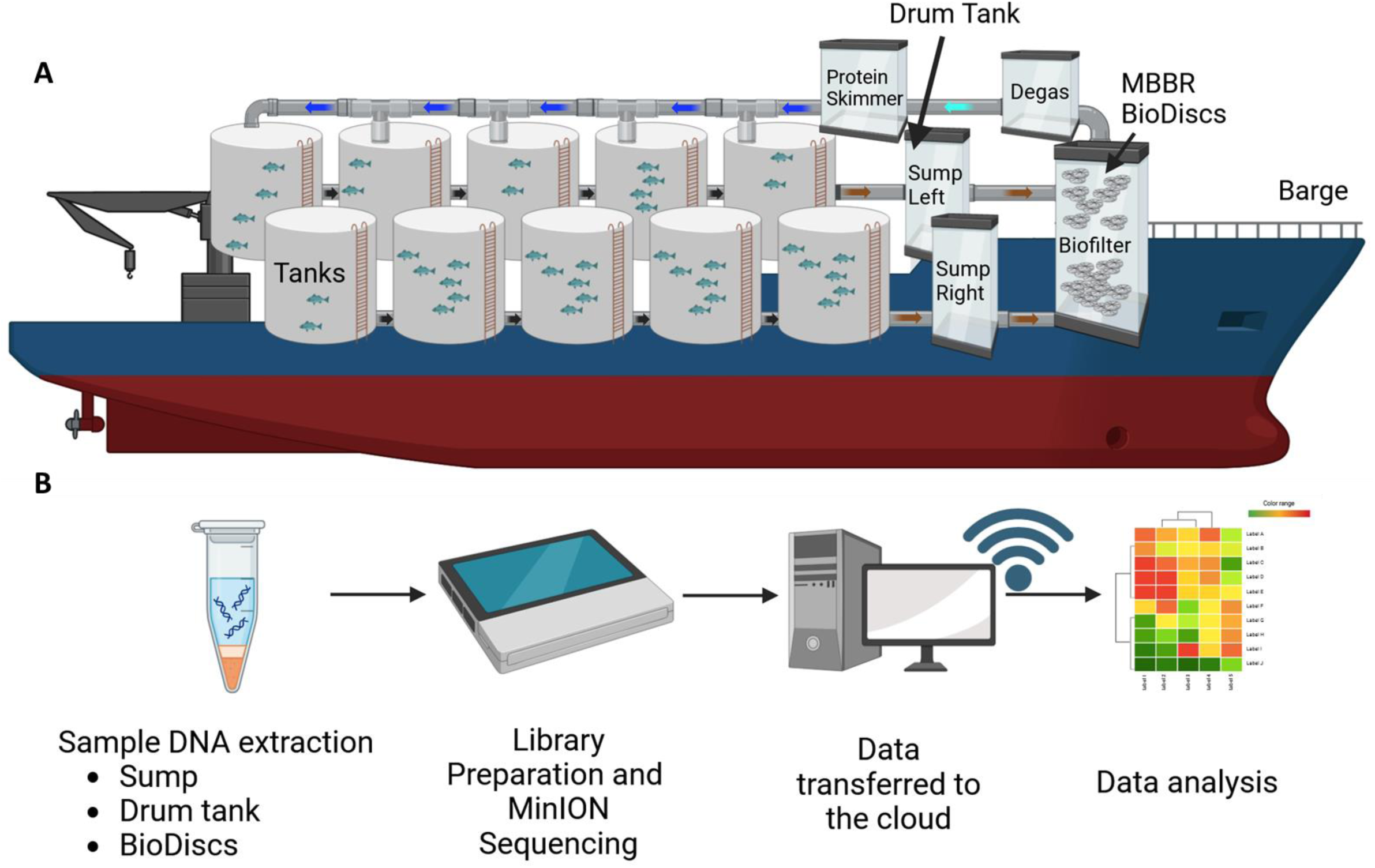
Diagram of the workflow from barge to detection. **A.** Cartoon of the barge used for the culture of fish species, in this case barramundi. The barge had 10 tanks holding up to 20,000 fish each. Samples were taken from the sides of the drum tank, from the floating BioDiscs and from the sump. **B.** The sample preparation workflow, starting from sample lysis to extraction of the DNA, library preparation using the rapid and rapid barcoding kits from Oxford Nanopore, followed by sequencing using the MinION device, upload of the data to a cloud server, whereupon data analysis is completed.

The aquaculture samples we have processed provide an insight into the profile of the aquaculture microbial community (Figure 2). A cutoff of 0.2% relative abundance from aligned bases was applied to provide insight into the taxa present at high to moderate relative abundance within the sump, drum tank and the BioDiscs. The presence of photosynthetic organisms was a curious finding as the facility is neither open to the sky nor lit with strong lighting, especially in the sump. However, organisms like *R. capsulatus* are capable of growth under low light conditions (80 lux) and are capable of fixing nitrogen, which could explain their presence.^17^ Taxa that thrive in saltwater were observed, *Muricauda ruestingensis* was first isolated from seawater sediment, which could explain why we observe the organism in high abundance in the sump and drum tanks; it had a mean relative abundance that is consistent in the sump and drum tank, and nearly all BioDiscs.^18^

**Figure 2.**
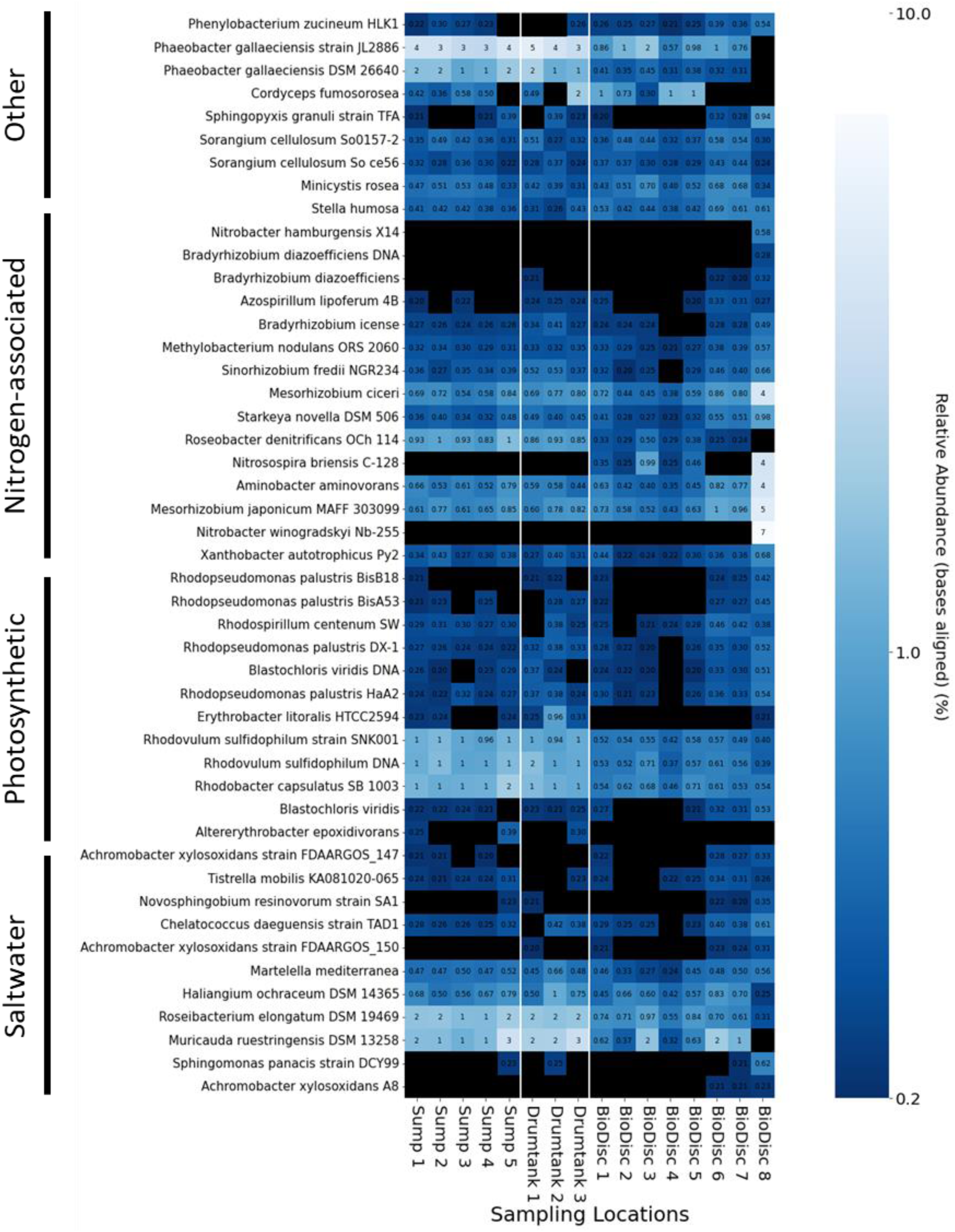
Aligned base relative abundance across the sampled aquaculture locations highlighting known broad categories of taxa. Aquaculture microbiome by sampling source, minimum N=3. Shown are the relative abundance as calculated by taking hitLength from centrifuge for each taxon and dividing by hitLength sum to get aligned bases for a given sample. Cut-off used was 0.2% relative abundance.

The set of organisms that we will focus upon for the remainder of this analysis will be literature identified nitrogen-associated organisms as they are a global concern to aquatic ecosystems and nitrogen compounds can negatively impact livestock through eutrophication and high ammonia.^19,20^ We examined the inter-sample diversity at the locations and found that variation within the samples appeared to cluster based on the location (Figure 3A). This suggests that the metagenomes identified were similar at the same sampling locations. Regarding intra-sample diversity, we observed that in the three sampling locations within the samples that alpha diversity was similarly rich, with the sump having the least evidence within the assessed samples (Figure 3B). We extracted reads from identified nitrogen-associated organisms, which are shown in Figure 3C. For each location, three biological replicates are depicted in the heat map. BioDiscs are partially submerged in wastewater, and provide a surface for microorganisms to grow and form a biofilm. The BioDisc rotates, the biofilm comes into contact with the tank water and air alternately.

**Figure 3.**
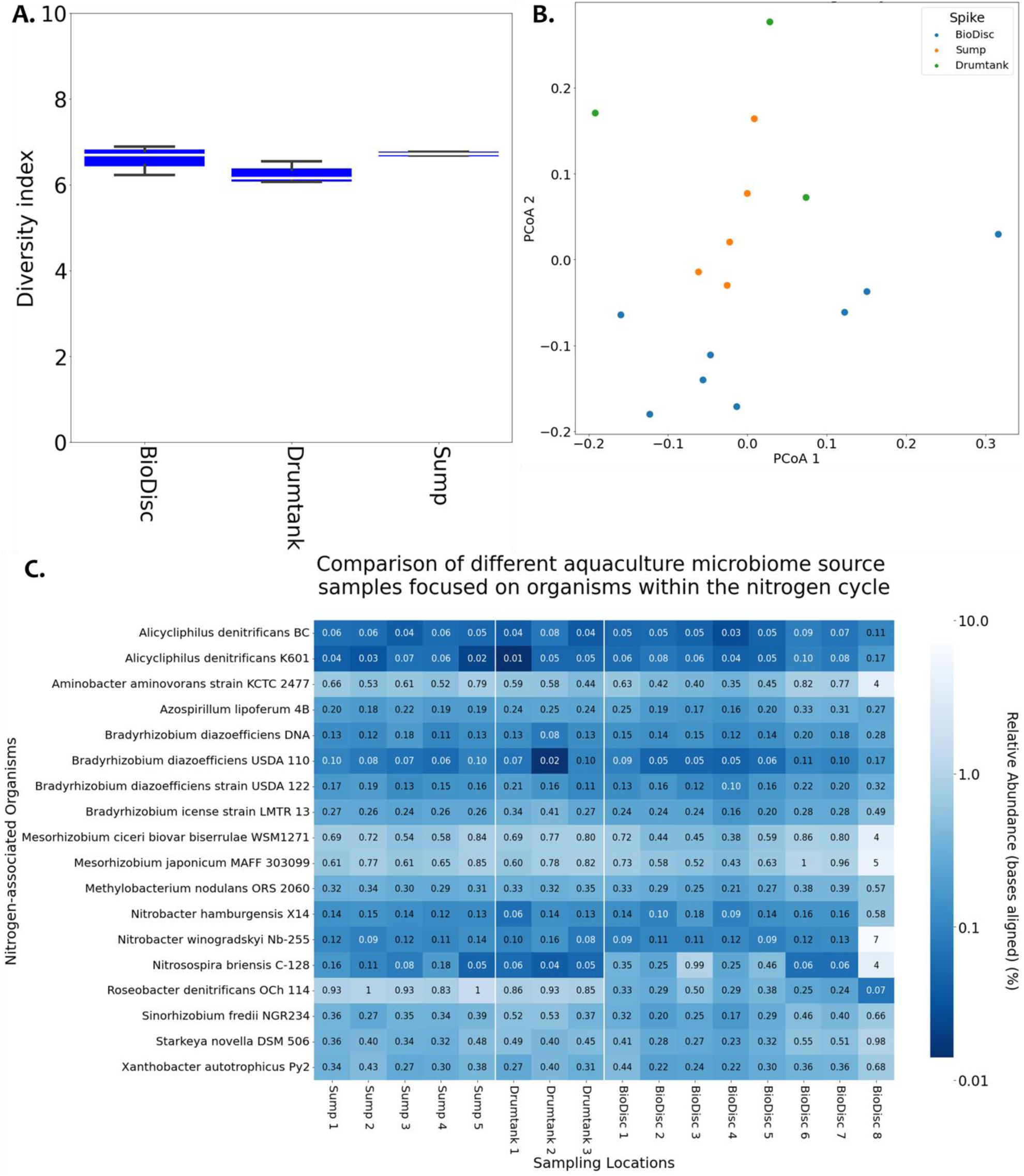
Baseline, nitrogen-associated microbiomes within the sampled aquaculture locations. **A.** Shannon alpha diversity. The three sampling locations are shown with the intra-sample diversity and the variation observed across the three samples. No significant difference in intra-sample diversity was seen between sample locations, Kruskal-Wallis H Test (H-value=4.3, p-value=0.11). **B**. Beta diversity analysis using principal coordinates analysis (PCoA) of the Bray-Curtis dissimilarity matrix (Eigenvectors: PC1: 37.98%, PC2: 25.41%). Multivariate Analysis of Variance (MANOVA) suggests that microbial composition varies systematically among the three sampling locations (Wilk’s Lambda, p=0.027). This is further corroborated by Permutational Multivariate Analysis of Variance (PERMANOVA), which for sampling location gave P=0.007, R^2^=0.302, F-value: 2.81. **C.** Aquaculture microbiome by sampling source, minimum N=3.

The trend that we observe is BioDiscs have the highest associated read count for nitrogen-associated organisms. This nitrogen-associated organism concentration decreases as we proceed away from the air-water interface and the BioDiscs. Potentially this could be related to biofilm supporting availability with more reads found for nitrogen-associated organisms in the sump than the drum tank (Figure 3C). We observed that in one instance, the BioDisc contained between 5,000 and 22,000 reads for *Nitrobacter* and *Nitrospira* species, while these organisms were not detected in other samples. However, we observed that for BioDisc-3, the overall read count for each nitrogen-associated organism is elevated relative to other samples.

## Discussion

Our workflow provides a tool for aquaculture metagenome monitoring, which is helpful to understand and react to growth tank functionality and infection events. The results clearly show that we are able to monitor the microbiome of the aquaculture tanks for nitrogen-associated organisms and other groups, including saltwater focused taxa within the three locations sampled. In the case of *Lates calcarifer* (barramundi), to prevent cannibalism, solutions including *ad libitum* feeding, stocking density and L-tryptophan supplementation were employed by the aquaculture staff. Excessive feeding can lead to nitrogenous waste and deteriorating water quality. Monitoring this parameter could allow for selection of a mix of probiotic nitrogen-associated organisms to mediate the water quality^21^ while recording probiotic efficacy and longevity in the recirculating system.

The selective presence of *Alicycliphilus denitrificans*, *Nitrobacter* and *Nitrospira* in one BioDisc could be influenced by the specific environmental conditions within the particular BioDisc, favouring the growth and activity of these nitrogen-converting bacteria. It’s plausible that BioDiscs provides optimal conditions for nitrification, with ideal oxygen, pH, and nutrient levels for *Nitrobacter* and *Nitrospira*, as compared to some other BioDiscs. It could also signify a higher efficiency in nitrogen conversion in this BioDisc, aiding in maintaining the nitrogen balance crucial for wastewater treatment. Understanding the exact parameters leading to this localised presence can offer insights for enhancing the performance of other BioDiscs by tweaking their environmental conditions to mimic this successful environment. Additionally, this observation underpins the necessity for consistent and detailed monitoring across different BioDiscs to ensure uniform and optimal performance in nitrogen management, essential for effective wastewater treatment and subsequent biomass monitoring. Our data revealed the presence of *Starkeya* and *Aminobacter* spp in the aquaculture system, especially at high concentrations on the BioDiscs. This suggests active participation not only in the nitrogen cycle but also in the sulfur cycle, given that *Starkeya* spp has a well-documented ability to oxidize thiosulfate. The thriving of *Starkeya* spp might hint at fluctuating nutrient conditions within the system, which would be conducive for organisms dependent on both sulfur and nitrogen compounds for metabolism. *Aminobacter*, known for its metabolic versatility, can engage in dissimilatory nitrate reduction to ammonia under certain conditions^22^. Additionally, some strains have demonstrated denitrification capabilities, converting nitrate and nitrite to nitrogen gas or nitrous oxide. While we observe high read counts for *Mesorhizobium* in the BioDiscs, we recommend caution as *Mesorhizobium* are known to have symbiotic relationships with leguminous plants to fix atmospheric nitrogen.

Working with metagenomic samples requires a careful and measured consideration for sample purity. This means that at multiple stages during the processing of these samples, we must be aware that noise can be introduced, such as from the DNA extraction, library preparation, as well as noise from the computational analyses. This is especially true when you are working with low-abundance samples. In this study, we worked with relatively high-abundance DNA inputs, however, to identify organisms that may be present at low concentrations, quality control measures would be required. In future experiments, we would include negative controls at the sample level, as well as a blank control for the DNA extraction step. This is important to catch deleterious contaminations to the barramundi culture but also to minimize the potential for misdiagnosis of an infection within the aquaculture system. Caution needs to be taken towards known false positive organisms which, according to the literature, include members of the *Bradyrhizobium* genus primarily, and also to a lesser extent in some cases, the *Mesorhizobium* genera^23–25^.

Other considerations that could be worked with here are the use of 16S and 18S rRNA gene protocols alongside the metagenomics analysis. We did not pursue an amplicon approach because our aim was to enable the aquaculture company to conduct these complex sequencing analyses directly at the aquaculture facility, and in a rapid manner. Incorporating tools like a thermocycler, for specific gene amplification, would unduly complicate matters and substantially increase the time requirements for this protocol. By minimizing the number of intricate steps, we sought to streamline the process, ensuring it remains both feasible and reproducible under near real-time, on-site conditions. Considerations for future *in-silico* bacterial detection using a 16S rRNA gene approach might consider using the MIDAS-4^26^ ribosomal DNA database, which has a comprehensive set of data for organisms extracted from wastewater treatment plants and may be relevant for aquaculture surveillance.

Application of the protocol enunciated in this study would enable aquaculture operations with responsive metagenome monitoring and health management. These processes would result in reduced transmission and incidence of disease, allowing consistently higher yields, as well as opportunities for optimization of the metagenomes.

## Acknowledgments

Funding for this research was provide by the Singapore National Biofilm Consortium (Grant SNBC/2021/ SF3/P02). In addition, this research was supported by the National Research Foundation, Prime Minister’s Office, Singapore under its Campus for Research Excellence and Technological Enterprise (CREATE) programme, through Singapore MIT Alliance for Research and Technology (SMART): Critical Analytics for Manufacturing Personalised-Medicine (CAMP) Inter-Disciplinary Research Group. Financial support from National Research Foundation and Ministry of Education Singapore under its Research Centre of Excellence Program was provided through Singapore Centre for Environmental Life Sciences Engineering.

## Author Contributions

JS: Conceptualization, Methodology, Software, Validation, Formal analysis, Data Curation, Writing - Original Draft, Writing - Review & Editing, Visualization. IB: Validation, Investigation, Resources, Writing - Review & Editing, Supervision. HM: Validation, Investigation, Resources. SS: Conceptualization, Resources, Writing - Original Draft, Writing - Review & Editing, Supervision, Project administration, Funding acquisition.

## Data Availability

Sequence Read Archive (SRA) ID **TBC**

